# Impact of a defined bacterial community including and excluding *Megamonas hypermegale* on broiler cecal microbiota and resistance to *Salmonella* infection

**DOI:** 10.1101/2024.08.15.608135

**Authors:** Camila Schultz Marcolla, Tingting Ju, Kimberlee Ten, Usha Sivakumar Sharma, Leakhena Moeun, Benjamin P. Willing

## Abstract

Intensive broiler production practices impair the transmission of commensal microbes from hens to offspring, resulting in a lower abundance of non-spore-forming strict anaerobic bacteria. We evaluated the effects of colonization by a defined community (DC) of bacteria including and excluding *Megamonas hypermegale* in chicks challenged with *Salmonella*. Inoculation with DC resulted in higher phylogenetic diversity and the dominance of Bacteroidetes species in the cecal microbiota, with a decrease in the relative abundance of *Salmonella* and *Escherichia/Shigella*, as well as a lower *Enterobacteriaceae* load. Substantial shifts in microbiota composition were coupled with subtle changes in metabolites and host responses, including changes in interferon-γ, macrophage colony-stimulating factor, propionate, valerate, and isovalerate concentrations in the ceca. We identified bacterial species that were able to establish and persist after a single exposure, many of which were members of Bacteroidetes. Although co-culture with *M. hypermegale* reduced *Salmonella* counts by 99.3% *in vitro*, *in vivo* inoculation of *M. hypermegale* increased splenic *Salmonella* counts in inoculated chicks. The use of DC containing bacteria isolates harvested from the cecal contents of mature chickens can recapitulate the changes in volatile fatty acid concentrations observed in birds colonized with complex communities, and the presence of *M. hypermegale* specifically enhances the production of propionate. Our findings suggest that the use of DC can be explored as a strategy to control disease occurrence in broiler production; however, further research is warranted to properly understand the role of individual species in the broiler cecal community aiding the formulation of appropriate DCs.

**Importance:** Intensive production practices can reduce beneficial gut bacteria in broiler chickens, potentially leading to higher disease risk. We investigated whether introducing a defined community (DC) of beneficial bacteria, along with *M. hypermegale*, could improve gut health and resistance to *Salmonella* in broiler chicks. Our findings show that DC increases microbial diversity and reduces the relative abundance of potential pathogens, like *Salmonella* and *Escherichia/Shigella*, which was coupled with subtle changes in the immune responses of the birds and higher concentration of volatile fatty acids in the ceca. This study suggests that using DC can enhance poultry health and reduce disease, providing a potential strategy to improve broiler production. However, further research is needed to understand the roles of individual bacteria and refine these bacterial communities for practical use in farming. This work holds promise for developing natural methods to enhance poultry health and safety.

## Introduction

Current poultry production practices aim to minimize bird exposure to pathogens that can cause disease and contaminate food products. These practices may impair the colonization of the chicken gastrointestinal tract with host-adapted commensal bacteria that have co-evolved with chickens in nature (Hird, 2017; Marcolla et al., 2023a; Nurmi and Rantala, 1973; Rantala and Nurmi, 1973). In fact, the cecal microbiota of broilers reared in intensive systems was shown to be less diverse and depleted of non-spore-forming strict anaerobic bacteria, especially from the phylum Bacteroidetes, compared to that of age-matched broilers from extensive systems (Marcolla et al., 2023a). The cecal microbiota of intensively raised broilers was shown to lack bacterial species that are present in extensively raised birds, including members of *Olsenella*, *Alistipes*, *Phocaeicola, Bacteroides*, *Barnesiella*, *Parabacteroides, Megamonas*, and *Parasutterella* genera (Marcolla et al. 2023a, Marcolla et al., 2023b; Ocejo et al., 2019; Ramírez et al., 2020; Seidlerova et al., 2020). It was also demonstrated that chicks inoculated with cecal contents and bacterial cultures derived from cecal contents were consistently colonized by *Alistipes*, *Phocaeicola, Bacteroides*, *Barnesiella*, *Mediterranea*, *Megamonas*, *Parabacteroides*, *Phascolarctobacterium*, and *Subdoligranulum*, indicating that these bacteria are highly adapted and able to colonize the chicken gut after a single early life exposure (Marcolla, et al. 2023b).

Species from the *Megamonas* genus, including *Megamonas rupellensis*, *Megamonas funiformis*, and *Megamonas hypermegale*, have been isolated from chickens (Medvecky et al., 2018; Poudel et al., 2022; Sakon et al., 2008) and were shown to be enriched in the gut microbiota of wild and free-range compared to intensively raised and domesticated birds (Ocejo et al., 2019; Scupham et al., 2008; Wienemann et al., 2011; Yadav et al., 2021). *M. hypermegale* is an anaerobic, gram-negative, non-spore-forming, and non-motile rod that was first isolated from turkey feces (Harrison and Hansen, 1963). It was shown to produce acetic, propionic, lactic, and trace amounts of succinic acids in broth culture (Barnes et al., 1979; Harrison and Hansen, 1963). Metagenomics analysis indicated that *M. hypermegale* can metabolize hydrogen, potentially reducing the accumulation of H_2_ that can hinder short-chain fatty acid production within the gut (Sergeant et al., 2014). *M. hypermegale* has also been correlated with *Salmonella* growth. One study reported reduction of *in vitro* growth of *Salmonella* Typhimurium by cross-streaking with *M. hypermegale;* however, the same study had shown no inhibition of *Salmonella* load in chicks inoculated with *M. hypermegale* at hatch and challenged with *S.* Typhimurium (Barnes et al., 1979). Nonetheless, chicks inoculated with a defined microbial community containing *Phocaeicola vulgatus* (formerly *Bacteroides vulgatus*) and *M. hypermegale* together with other 46 bacterial isolates were shown to have higher resistance to *S.* Typhimurium infection (Impey et al., 1982b). Importantly, when chicks were inoculated with the same defined microbial community and received a diet containing antibiotics depleting *P. vulgatus* and *M. hypermegale*, no inhibitory effect of the defined microbial community on *Salmonella* levels was observed, suggesting that *P. vulgatus* and *M. hypermegale* may play a key role in host resistance to *Salmonella* infection (Impey et al., 1982a).

In addition to the association between *M. hypermegale* and *Salmonella*, *M. hypermegale* abundance has also been negatively associated with *Campylobacter jejuni* abundance in the turkey gut (Scupham et al., 2010). Broilers receiving cecal microbiota transplant showed an increase in *M. hypermegale* abundance associated with reduced severity of necrotic enteritis symptoms (Zaytsoff et al., 2022). Since the evidence collectively suggested the importance of *M. hypermegale* as a core member of the poultry gut microbiome, we hypothesized that this commensal would effectively engraft and affect bird physiology once introduced to newly hatched chicks. The current study aimed to evaluate the effect of early-life introduction of *M. hypermegale* alone or in combination with a defined community (DC) of bacteria on broiler gut microbiota development, host immune responses, and ability to resist *Salmonella* infection.

## Materials and methods Ethics approval

The animals used in this study were housed and maintained according to requirements of Canadian Council on Animal Care and Canadian Biosafety Standards for Facilities Handling or Storing Human and Terrestrial Animal Pathogens and Toxins. This study was approved by the University of Alberta Animal Care and Use Committee (AUP00002572 and AUP00001626).

### Preparation of microbial inocula

To isolate commensal bacteria, cecal digesta were collected from broilers and layers raised on commercial farms across Alberta. Isolation, culturing, identification, and sequencing procedures were described previously (Marcolla et al., 2023a). Species to be incorporated into the DC were selected based on the ability of isolates to grow easily in the lab, their abundance within the broiler cecal microbiota, and their ability to colonize the chicken gut according to our previous findings (Marcolla et al., 2023a; Marcolla, et al., 2023b). The DC was comprised of selected isolates including the following species: *Alistipes finegoldii*, *Bacteroides meditarraneensis*, *P. vulgatus*, *Fournierella massiliensis*, *Ligilactobacillus agilis*, *Ligilactobacillus aviarius*, *Lactobacillus crispatus*, and *Subdoligranulum variabile*. Four novel isolates, to each the nearest matches according to orthoANI values after whole genome sequencing were *Ruminococcus torques* (74.3%*), Bacteroides gallinaceum* (74.2%)*, Bacteroides uniformis* (75.03%), and *Barnesiella viscericola* (85.73%) were also incorporated in the DC. In addition, *M. hypermegale* was added to the DC community for the treatment defined as DC + Mega. All isolates were cultured in fastidious anaerobe agar (Neogen, USA), except for *Lactobacillus* and *Ligilactobacillus* species, which were cultured on de Man Rogosa and Sharpe (MRS) agar (BD Difco, USA). Cultures were incubated under anaerobic conditions (5% CO_2_, 5% H_2_, and 90% N_2_) in an anaerobic chamber (Bactron300, Sheldon Manufacturing, USA) for 48 h at 37°C. Colonies were picked and re-inoculated into fastidious anaerobe broth (FAB) or MRS broth. Bacterial cells collected from the broth culture or agar plates were mixed with 50% glycerol at 1:1 volume, aliquoted into 1.5 ml tubes to make glycerol stocks and stored at -80°C. Prior to being inoculated to birds, glycerol stocks of each isolate were thawed on ice in an anaerobic chamber and equal volumes of each isolate were pooled. Fresh *M. hypermegale* cultured in FAB was added to the DC for the DC + Mega treatment.

*Salmonella enterica* serovar Enteriditis SGSC 4901 (PT4) and *Salmonella enterica* serovar Typhimurium ATCC SL1344 were streaked on fresh xylose-lysine-deoxycholate agar (XLD) (Thermo Scientific) and incubated for 18 h at 37 °C. A single colony from each plate was selected, inoculated in 5 ml FAB, and incubated for 18 h at 37 °C. After incubation, broth culture was serial diluted in sterile 1 x phosphate buffered saline, which was subsequently plated onto XLD plates and incubated for 18 h at 37 °C for bacterial enumeration. The broth was diluted to achieve 1 x 10^8^ colony forming units (CFU)/ml.

### Acids, bile, and oxygen tolerance of M. hypermegale and Salmonella inhibition in vitro

To evaluate the tolerance of *M. hypermegale* to acids, bile and oxygen, *M. hypermegale* was seeded into 5 ml of brain heart infusion broth (BHI, Oxoid, CA) and incubated at 37°C for 48 h. For acid tolerance assay, 500 μl of the seeded broth was inoculated into 4.5 ml of BHI broth adjusted to pH 2, 3, 5, and 7. For bile tolerance assay, 500 μl of the seeded broth was inoculated into BHI broth containing 0.3%, 0.6%, and 1.2% of sterile porcine bile. All samples were incubated anaerobically at 37°C for 3 h determined based on the average transit time of digesta through the chicken gizzard and small intestine (Svihus and Itani, 2019). For the oxygen tolerance assay, 500 μl of the seeded broth was inoculated into 4.5 ml of BHI broth and incubated aerobically at 37°C for 15, 30, and 60 min. A control sample not exposed to oxygen was considered as time 0. After treatments, samples were plated on BHI agar and incubated anaerobically at 37°C for 48 h.

*In vitro* assays were performed to investigate the ability of *M. hypermegale* to inhibit *S.* Typhimurium growth using co-culture and agar slab method. Broth cultures containing 10^4^ CFU/ml of *S.* Typhimurium or *M. hypermegale* were co-inoculated into BHI broth. A BHI broth inoculated with 10^4^ CFU/ml of *S.* Typhimurium was used as control. Samples were incubated anaerobically at 37°C for 48 h, then plated on XLD agar for *S.* Typhimurium enumeration. The inhibition effect was determined by comparing the *S.* Typhimurium load in the co-culture relative to the control sample. The inhibitory effect of *M. hypermegale* on *S.* Typhimurium was also tested using the “agar slab method” (Dec et al., 2014). Briefly, broth containing 10^4^ CFU/ml of *M. hypermegale* was spread onto BHI agar, incubated anaerobically at 37°C for 48 h, and agar slabs measuring 9 mm in diameter were cut and placed onto a BHI plate spread with *S.* Typhimurium, which was further incubated at 37°C for 24 h to measure the zone of inhibition.

### Animal housing and study design

Day-old broilers (Ross 708, Aviagen, Huntsville, AL) obtained from a commercial hatchery were weighed, tagged with individual IDs, and randomly distributed into two-level individually ventilated isolators (GR1800 double decker Sealsafe® plus, Tecniplast, CA) lined with sterile aspen shavings. Three chicks were housed in each cage with *ad libitum* access to water and food (Laboratory Chick Diet S-G 5065, LabDiet, MO, US) throughout the experiment. Cages were changed as needed and 50 g of bedding materials from the dirty cage were transferred to clean cages to promote exposures to seeded microorganisms. All procedures were performed in a biosafety cabinet. Cages were kept in a temperature-controlled room, with a daily lighting schedule of 12 h light. Room temperature was kept at 30°C for the first three days of age and then gradually reduced to 24°C as birds aged. At the beginning of each experiment, ten chicks were euthanized at arrival and cecal samples were collected and plated on XLD agar to confirm the absence of *Salmonella* in the baseline microbiome.

In a preliminary experiment (EXP1), 60 day-old chicks weighing 47.24 ± 7.25 g (mean ± standard deviation (SD)) were randomly distributed into isolators (3 birds/isolator) and allocated into four treatments: Control, Mega, DC, and DC + Mega. Control chicks were inoculated with sterile liquid casein yeast extract medium; chicks from the Mega treatment were inoculated with frozen *Μ. hypermegale* glycerol stock containing 5x10^5^ CFU/ml of cells; chicks from the DC treatment were inoculated with DC isolates, while chicks from the DC+Mega treatment were inoculated with DC isolates and *M. hypermegale* glycerol stock. All inoculations were performed the day after arrival via oral gavage with 150 μl of inocula.

Results from EXP1 indicated that *M. hypermegale* inoculated as a frozen glycerol stock failed to colonize the chicken gut. Thus, a follow-up experiment (EXP2) was designed to test the colonization ability of *M. hypermegale* when provided as a fresh broth culture. Specifically, a total of 48 day-old chicks weighing 44.9 ± 3.8 g (mean ± SD) were randomly distributed into cages and assigned to Control or Mega treatments. Chicks in the Control treatment were inoculated with sterile LCY, while chicks from the Mega treatment were inoculated with 150 μl of fresh *M. hypermegale* broth containing 5x10^5^ CFU/ml. Inoculations were performed at the day of arrival and repeated at 48 h after arrival. Two days after the repeated inoculation, chicks in all treatments were infected with *S.* Enteritidis by oral gavage with 1.5 x10^6^ cells/bird. On day 3 post infection (5 days of age), the lightest chick in each cage was selected for sampling, while the 2 remaining chicks were sampled 10 days post infection (13 days of age).

Results of EXP2 indicated that *M. hypermegale* successfully colonized the gut when introduced as a fresh broth culture, and a third experiment (EXP3) was conducted using fresh broth culture of *M. hypermegale* in combination with the DC. A total of 72 day-old chicks weighing an average of 46.7 ± 3.7 g (mean ± SD) were randomly distributed into twenty-four cages (3 birds per cage) and assigned to three treatments: Control, DC, and DC + Mega (8 cages per treatment). The day after arrival, control chicks were inoculated with sterile LCY; chicks in the DC treatment were inoculated with DC isolates; and chicks in the DC + Mega treatment were inoculated with DC isolates and fresh *M. hypermegale* broth (1x10^5^ CFU/ml). Inoculations were repeated 24 h later, and 48 h after the second inoculation all chicks were infected by oral gavage with 150 μl of 1x10^7^ CFU/ml *S.* Enteritidis PT4. The lightest chick in each cage was sampled 48 h after infection, whereas the two remaining chicks in each cage were sampled 9 days post-infection (13 days of age).

### Sampling

Chickens were euthanized by cervical dislocation, and the coelomic cavity was opened using sterile technique. The whole spleen was collected into 1 x phosphate buffered saline, stored on ice for transport, homogenized by bead-beating twice at 6.0 m/s for 40 s (FastPrep-24TM 5G, MP Biomedicals), plated on XLD and incubated for 24 h at 37°C for *Salmonella* enumeration. Cecal digesta and tissues were collected and immediately stored at -80°C. Concentrations of short-chain fatty acids (SCFA) in cecal digesta were determined by gas chromatography as described previously (Marcolla et al., 2023b). Levels of interferon (IFN)-α, ΙFN-γ, interleukin (IL)-2, IL-6, IL-10, IL-16, IL-21, macrophage colony-stimulating factor (M-CSF), macrophage inflammatory protein (MIP)-1β and MIP-3α, regulated on activation, normal T cell expressed and secreted (RANTES) chemokine, and vascular endothelial growth factor (VEGF) in cecal tissue were determined by a multiplex cytokine assay (Featured – Chicken Cytokine/ Chemokine 12-Plex Assay, Eve Technologies Corporation, CA) as described previously (Marcolla et al. 2023b). Ileal segments collected at 0.5 cm proximal and 0.5 distal to Meckel’s diverticulum were prepared for histomorphology analysis and stained with hematoxylin eosin as previously described (Marcolla et al., 2023a). Images of representative cross-sections of ileal samples were taken using BioTek Lionheart Imager FX (Agilent technologies) and analysed using Agilent Gen 5 software (Gen5 v.3.12).

### Salmonella, Enterobacteriaceae, and Bacteroidetes quantification

Quantitative RT-PCR was used to determine total bacteria, *Salmonella*, *Enterobacteriaceae*, and Bacteroidetes load in cecal contents. To generate standards curves, DNA was extracted from pure broth cultures of *S.* Enteritidis and *Alistipes finegoldii* using a Wizard Genomic DNA Purification Kit (Promega Corporation, WI, USA) following the manufacturers’ protocol. The *Salmonella* enterotoxin gene (*stn*) gene was amplified using forward primer, 5’-CTTTGGTCGTAAAATAAGGCG-3’ and reverse primer, 5’- TGCCCAAAGCAGAGAGATTC-3’ (Ziemer & Steadham, 2003). The quantification of Bacteroidetes was performed using forward primer, 5’-GGARCATGTGGTTTAATTCGATGAT -3’ and reverse primer, 5’- AGCTGACGACAACCATGCAG -3’ (Guo et al., 2008). The quantification of *Enterobacteriaceae*was performed using forward primer, 5’- CATTGACGTTACCCGCAGAAGAAGC -3’ and reverse primer, 5’- A CTCTACGAGACTCAAGCTTGC -3’ (Bartosch et al., 2004). Total bacteria were quantified using forward primer, 5’- TCCTACGGGAGGCAGCAGT -3’ and reverse primer, 5’- GGACTACCAGGGTATCTAATCCTGTT -3’(Nadkarni et al., 2002). The reaction mixtures contained 5 μl of SYBR Green SuperMix (Quantabio, US), 0.5 μl of each forward and reverse primer, 3 μl of nuclease free water, and 1 μl of DNA diluted to a concentration of 5 ng/μl. The PCR programs consisted of an initial denaturation step of 3 min at 95°C, followed by 40 cycles of 95°C for 10 s and 60°C for 30 s, which was performed on an ABI StepOne™ real-time System (Applied Biosystems, Foster City, CA).

### DNA extraction and 16S rRNA gene amplicon sequencing analysis

Total DNA from cecal digesta was extracted using a QIAamp DNA stool mini kit (Qiagen NV, Netherlands), following the manufacturers’ Pathogen Detection protocol, with minor modifications. Specifically, approximately 100 mg of digesta content was mixed with the Inhibitex® buffer and 2.0 mm garnet beads (BioSpec Products, Bartlesville, OK), homogenized and lysed by bead-beating twice at 6.0 m/s for 30 s. DNA concentrations were determined using Quant-iT™ Picogreen™ dsDNA assay kit (Invitrogen, Thermo Fisher Scientific, US).

Amplicon libraries targeting the V3-V4 regions of the *16S rRNA* gene were prepared following the Illumina 16S Metagenomic Sequencing Library Preparation protocol (#15044223 Rev.B). Sequencing was performed on an Illumina MiSeq platform (Illumina Inc, San Diego, CA) using 2x 300 cycles. Raw sequences were processed using Quantitative Insights into Microbial Ecology 2 v2020.2 (Bolyen et al., 2019). Forward and reverse sequences were denoised and truncated at 270 and 220 bp, respectively, and chimeras were removed using DADA2 (v. 2020.2.0) plugin (Callahan et al., 2016). Multiple sequence alignments were performed using MAFFT (Katoh & Standley, 2013) and phylogenetic trees were generated using FastTree method (Price et al., 2010). Naïve Bayes classifier (Pedregosa et al., 2011) pretrained on SILVA 138 QIIME compatible database (Quast et al., 2013) was used for taxonomic classification, and sequences were clustered at 99% identity using majority taxonomy strings. Downstream analyses were performed using phyloseq v.1.40.0 (McMurdie & Holmes, 2013), microbiome v.1.18.0 (Lahti & Shetty, n.d.) and qiime2R v.0.99.6 (Bisanz, n.d.) packages in R v 1.4.1717 (RStudio Team, 2020). Amplicon sequence variants (ASVs) assigned to Mitochondria family, Chloroplast order, Archaea kingdom, Unassigned at the phylum level and present in less than 10% of the samples or presenting less than 10 reads were removed from the dataset. Reads were rarefied at 22,006 reads before downstream analysis. Alpha-diversity was evaluated using phylogenetic diversity and Chao1 indices. Beta-diversity was evaluated using Bray-Curtis distance matrix and visualized by principal coordinates analysis (PCoA). Differentially abundant taxa were identified using DESeq2 with apeglm for logarithmic fold change shrinkage and FDR correction analysis (Love et al., 2014; Zhu et al., 2019). Analysis at the taxa level was performed by merging all the ASVs exhibiting the same taxonomy string using *tax_glom* function (phyloseq package). Spearman correlation analysis was performed using psych v.2.3.3 package (Revelle, 2023). Figures were generated using ggplot2 v.3.4.0 (Wickham, 2016).

### Colonization ability and efficiency

Colonization ability and colonization efficiency of bacterial isolates were determined based on the prevalence and relative abundance of each isolate in the cecal samples of inoculated birds. Specifically, bacteria that were detected in at least half of the inoculated birds were considered as prevalent, thus having high colonization ability. Colonization efficiency was determined by dividing the average number of reads of each isolate in the inoculated birds by the number of reads of that isolate in the inocula. Colonization efficiency results lower than 0.5, between 0.5 and 1, and higher than 1 were classified as low, medium, and high efficiency colonizers, respectively.

### Statistical analyses

All statistical analysis were performed in R (RStudio Team, 2020). Beta-diversity matrices were analysed using multivariate homogeneity of group dispersions and permutational multivariate analysis of variance with Benjamin-Hochberg procedure for FDR control using phyloseq v.1.40.0 (McMurdie & Holmes, 2013) and vegan v. 2.6-4 (Oksanen J et al., 2022) packages. Data were tested for normality using the Shapiro-Wilk test and analyzed by one-way ANOVA followed by Tukey’s HSD test if normally distributed, or Kruskal-Wallis and pairwise Dunn test with Bonferroni adjustment for multiple comparisons if distribution was not normal (stats package v. 4.3.3). A *p*-value of less than 0.05 was considered as statistically significant.

### Data availability

The 16S rRNA sequencing data generated and analyzed in the current study are available in NCBI SRA repository PRJNA1144928.

## Results

There were no differences in the body weight among treatments at any of the timepoints measured in all three trials (Table 1). Histological analyses performed on ileal tissues of birds in EXP3 indicated no differences in villus height, crypt depth, villus width and villus height:crypt depth ratios of birds in control, DC and DC + Mega treatments (Table 2).

**Table 1.** Body weight (average ± SD, g) in all treatments across experiments.

| Treatment | Control | Mega | DC | DC + Mega |
| --- | --- | --- | --- | --- |
| **14-days-old** |  |  |  |  |
| EXP1 | 409.5 ± 47.1 | 430.2 ± 61.5 | 415.7 ± 47.3 | 419.3 ± 48.7 |
| EXP2 | 413.5 ± 52.2 | 408.9 ± 61.8 | NA | NA |
| EXP3 | 367.5 ± 32.2 | NA | 368.1 ± 51.4 | 363.6 ± 49.2 |
| **7-days-old** |  |  |  |  |
| EXP1 | 144.9 ± 14.1 | 153.9 ± 13.9 | 152.1 ± 18.3 | 154.7 ± 19.9 |
| EXP2 | 177.7 ± 25.5 | 183 ± 23.4 | NA | NA |
| EXP3 | 137.6 ± 11.0 | NA | 139.9 ± 51.4 | 138.6 ± 14.8 |
| **1-day-old** |  |  |  |  |
| EXP1 | 48.6 ± 4.0 | 48.4 ± 5.9 | 48.5 ± 3.5 | 48.4 ± 5.6 |
| EXP2 | 45.6 ± 3.3 | 46.0 ± 3.2 | | |
| EXP3 | 47.4 ± 3.1 | NA | 47.2 ± 3.0 | 47.7 ± 3.3 |

**Table 2.** Effect of control, DC, and DC + Mega treatments on ileum morphology of 14-day- old broilers in experiment 3^a^.

|  | VH | SEM | CD | SEM | VW | SEM | VH/CD | SEM |
| --- | --- | --- | --- | --- | --- | --- | --- | --- |
| Control | 599.42 | 13.34 | 114.82 | 3.60 | 81.47 | 2.83 | 5.53 | 0.18 |
| DC | 665.64 | 18.68 | 128.05 | 4.41 | 85.42 | 3.42 | 5.47 | 0.21 |
| DC + Mega | 628.15 | 10.21 | 121.90 | 4.47 | 82.43 | 2.90 | 5.50 | 0.20 |
| *p*-value<sup>b</sup> | 0.095 |  | 0.72 |  | 0.15 |  | 0.91 |  |

<sup>a</sup>VH, mean villus height (μm), CD, mean crypt depth (μm); VW, mean villus width (μm), VH/CD, mean villus height to crypt depth ratio; SEM, standard error of the mean.
<sup>b</sup> Means were compared using Kruskal-Wallis test.

### Salmonella growth was inhibited by co-culturing with M. hypermegale in vitro

*M. hypermegale* can endure pH 5, at least 30-min of oxygen exposure, and up to 1.2% bile acid in the media. There was no difference in viable cells between pH 5 and 7 (*p* = 0.943), while no survival was observed at pH 2 or 3. The presence of up to 1.2% of bile in the media had no effects on *M. hypermegale* survival (*p* = 0.374). Oxygen exposure for 60 min reduced the number of viable cells of *M. hypermegale* (*p* = 0.047) by 41.7%, but no effects on survival were observed after 15- and 30-min of oxygen exposure compared to initial bacterial abundance (time 0). Co-culture of *M. hypermegale* and *Salmonella* reduced *Salmonella* counts by 99.3% (*p* = 0.003); however, no inhibition zone was observed using the agar slab method.

### M. hypermegale introduced as fresh culture was an efficient colonizer

In EXP1, *M. hypermegale* introduced from a frozen glycerol stock failed to colonize the chicken ceca and was not detected in the Mega or DC + Mega birds. At day 14, beta-diversity analysis indicated that the cecal microbiota of Control and Mega birds was different from that of DC and DC + Mega birds (r^2^=0.70, *p* = 0.001); however, no differences were found in the cecal microbiota between Control and Mega birds, nor between DC and DC + Mega birds (Figure 1A). The DC + Mega birds presented higher PD than that of Control (*p* = 0.03) and Mega (*p* = 0.01) birds, but no differences were observed in Chao1 index (*p = 0.26*) (Figure 2A). In EXP2, *M. hypermegale* introduced from a fresh culture successfully colonized the chicken gut. Gut microbial structures, as indicated by beta-diversity matrix, were different between Control and Mega birds (r^2^ = 0.60, *p* = 0.001) (Figure 1B). No differences in PD and Chao1 index (*p* = 0.355, *p = 0.96*, respectively) were found between Control and Mega birds (Figure 2B). In EXP3, beta-diversity analysis indicated that the cecal microbiota of control group was different from that of DC and DC + Mega (r^2^ = 0.64, *p* = 0.001 and r^2^ = 0.71 and *p* = 0.001, respectively), and that DC and DC + Mega microbial communities also differed (r^2^ = 0.26, *p* = 0.001) (Figure 1C). The microbiota of birds treated with DC or DC + Mega presented higher PD than the Control group (*p* < 0.001), and Chao1 index was higher in DC + Mega compared to Control group (*p* < 0.001) (Figure 2C).

**Figure 1.**
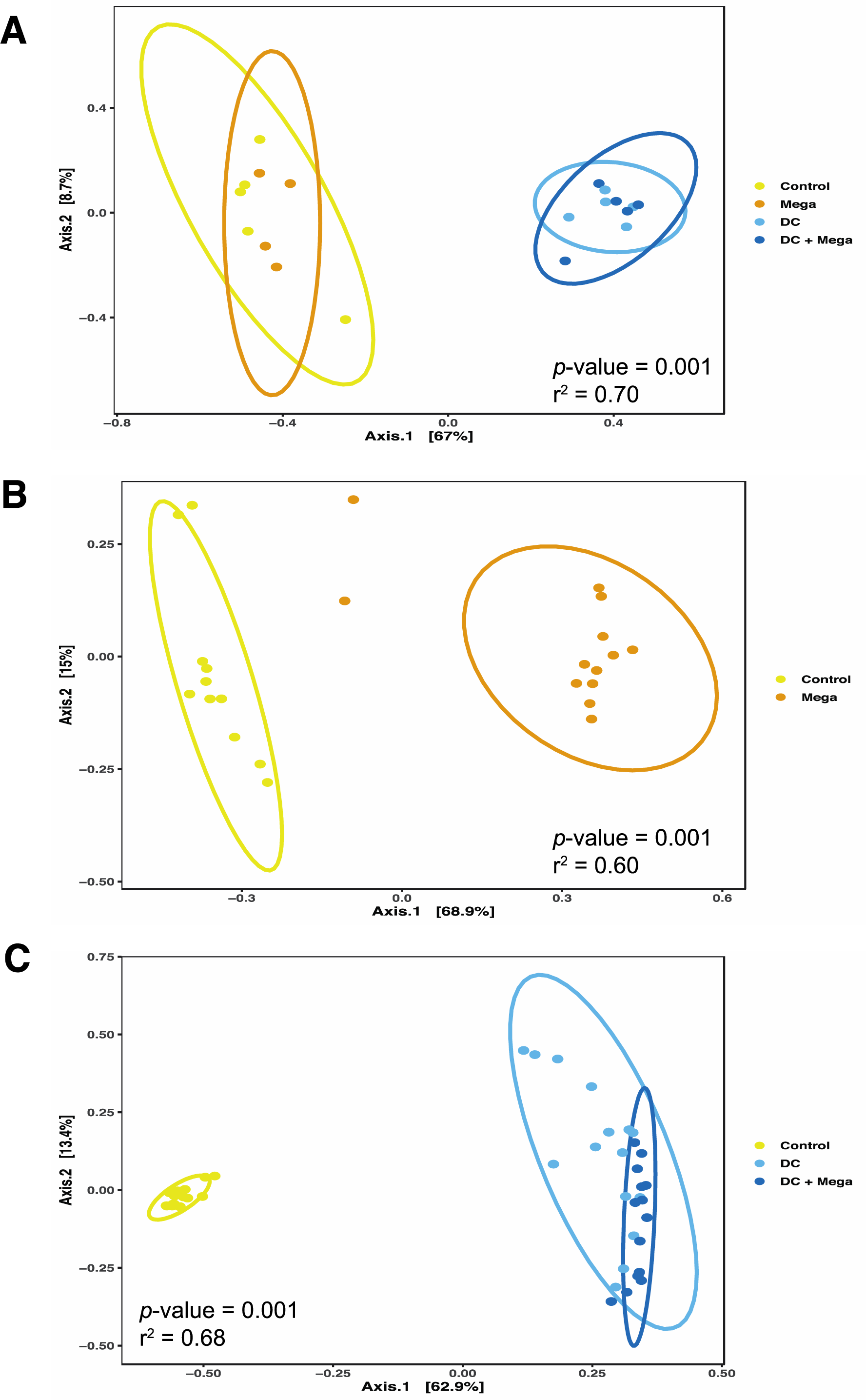
Effect of inocula on microbial beta-diversity. Principal coordinates analysis (PCoA) generated based on Bray-Curtis dissimilarity matrix of cecal samples obtained from 14-day-old Control chicks and chicks colonized with Mega, DC or DC + Mega treatments in EXP1 (A), EXP2 (B), and EXP3 (C). Samples are colored and shaped according to treatments and data ellipses represent the 95% confidence region for group clusters assuming a multivariate t-distribution.

**Figure 2.**
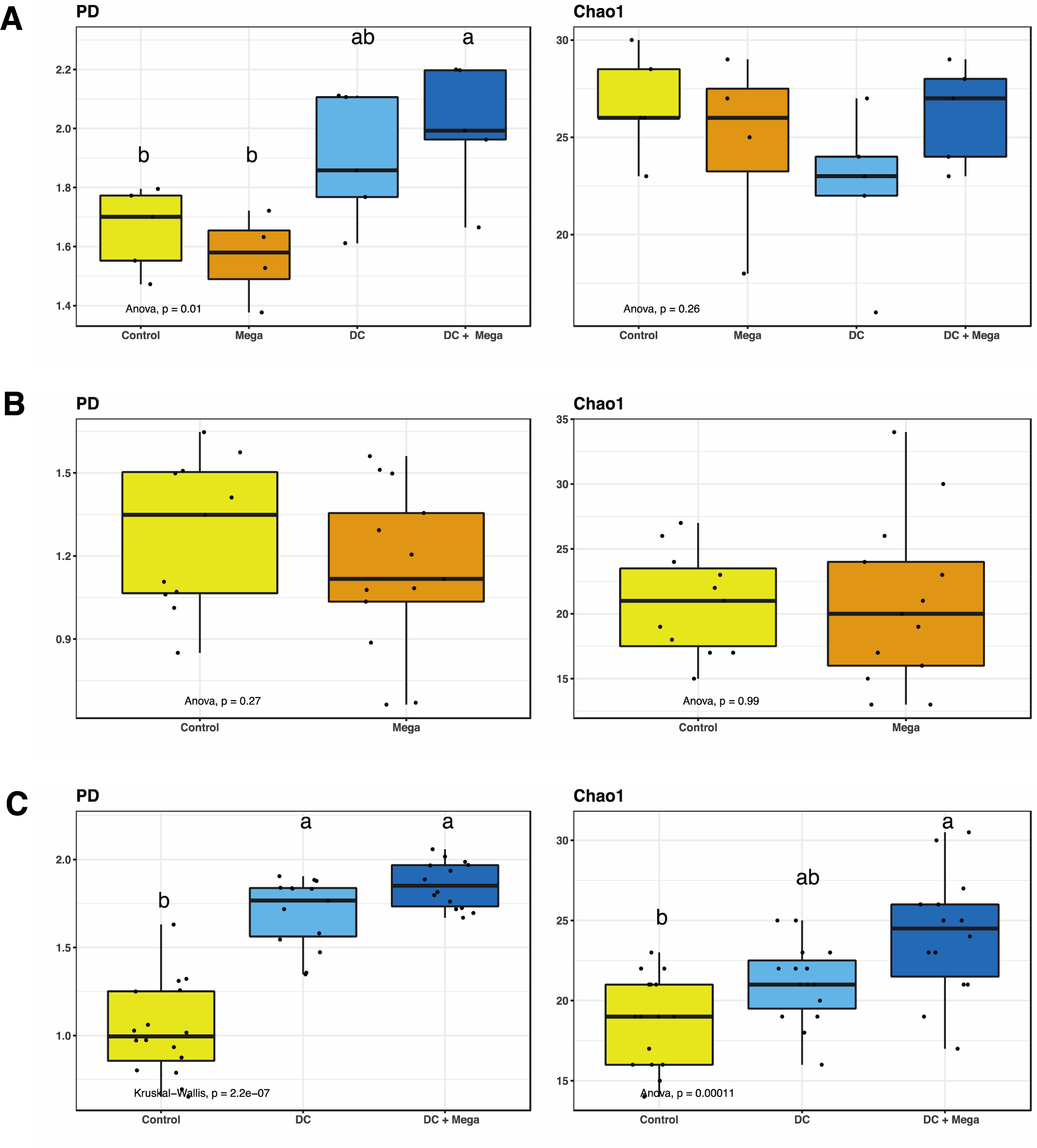
Alpha-diversity indices PD and Chao1 of cecal samples obtained from 14-day-old Control chicks and chicks colonized with Mega, DC or DC + Mega treatments in EXP1 (A), EXP2 (B) and EXP3. Superscripts with different letters indicate significant differences at α = 0.05.

In EXP1 and EXP3, an average of 94.0 ± 4.0 % of the cecal microbial community of DC and DC + Mega groups were bacteria found in the DC inoculum, whereas only 15.2 ± 18.5% of the cecal microbial community in Control birds was shared with the DC inoculum. Bacteria shared between Control and DC inocula included: *F. massiliensis* detected in one control bird in EXP1 at 60.9% relative abundance; *L. agilis* detected in three out of 16 birds in EXP3 with average relative abundance of 28.8 ± 20.5%, and *L. crispatus*, which was consistently detected in all Control birds from EXP3, with average relative abundance of 10.7 ± 12.7%. Due to consistently being detected in all control birds, *L. crispatus* was considered as a shared member of the baseline/initial microbiota in EXP3 (Figure 3). In EXP3, the cecal microbiota of Control birds was dominated by Firmicutes (an average of 50.8 ± 12.8 %) and Proteobacteria (an average of 49.2 ± 12.8 %), whereas the cecal microbiota of DC and DC + Mega birds were dominated by Bacteroidetes (an average of 86.3 ± 7.1% and 73.1 ± 6.3 %, respectively), and Firmicutes (an average of 8.8 ± 5.8 % and 24.8 ± 6.3 %, respectively) and Proteobacteria (an average of 4.8 ± 2.5 % and 2.0 ± 0.9 %, respectively) made up a small proportion of the community. Consistently, qPCR results indicated birds in DC and DC + Mega treatments to have a higher load of Bacteroidetes (*p* < 0.001) and a lower load of *Enterobacteriaceae* (*p* = 0.002) than the birds in control groups, while no differences in total bacterial load were detected (*p* = 0.58).

**Figure 3.**
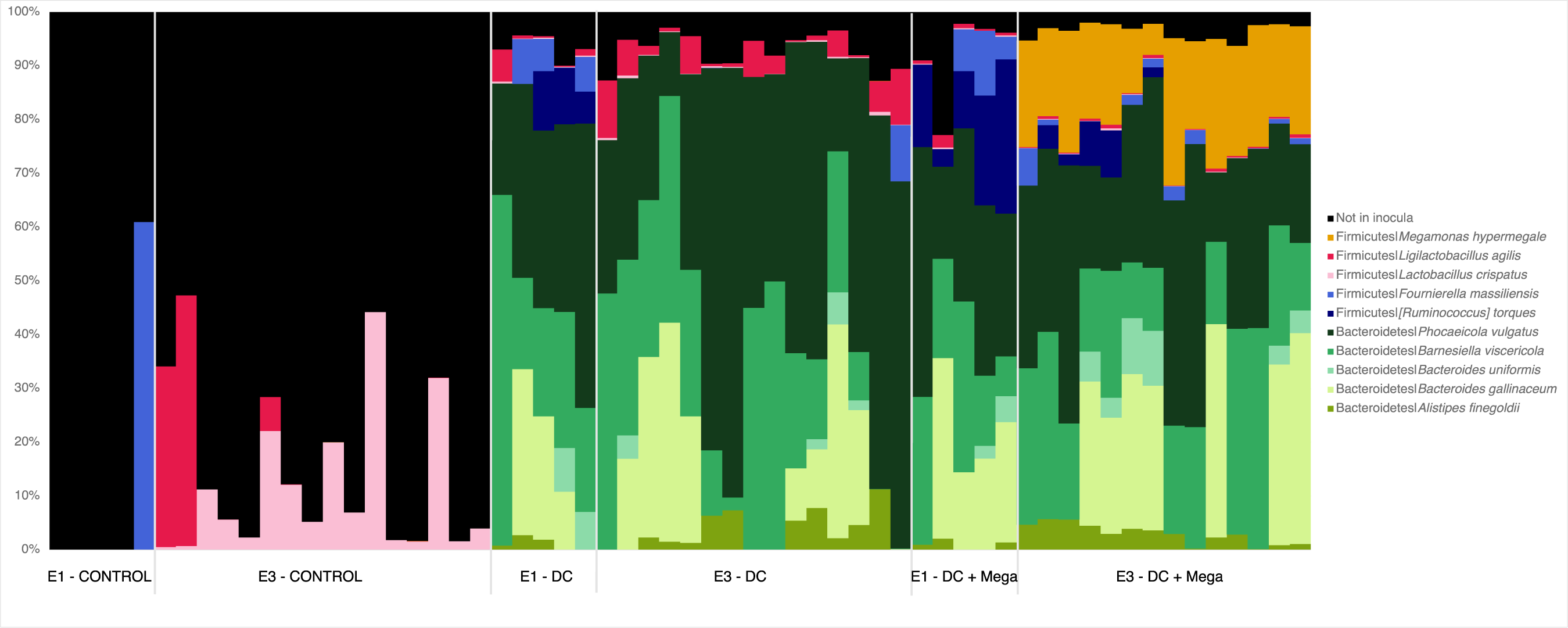
Bar plots of the relative abundances of bacterial species included in the DC and DC + Mega inocula detected in cecal samples obtained from 14-day-old chicks from Control, DC, and DC + Mega treatments in EXP1 and EXP3. Species not included in the inocula were considered as baseline microbiota and combined into as “Not in inocula” and shown in black.

In EXP2, *M. hypermegale* given as a fresh culture successfully colonized the ceca of treated birds and relative abundance ranged from 18% to 73%, with an average of 57.0 ± 15.9% (Figure 4A). *M. hypermegale*-treated birds presented lower relative abundance of *Enterococcus* (*p* = 0.001) and *Escherichia-Shigella* than Control birds (*p* < 0.001). There were no differences in cecal *Salmonella* load as determined by qPCR (Figure 4B); however, enumeration technique showed a higher *Salmonella* load in the spleen of Mega-treated birds (*p* = 0.002, Figure 4C).

**Figure 4.**
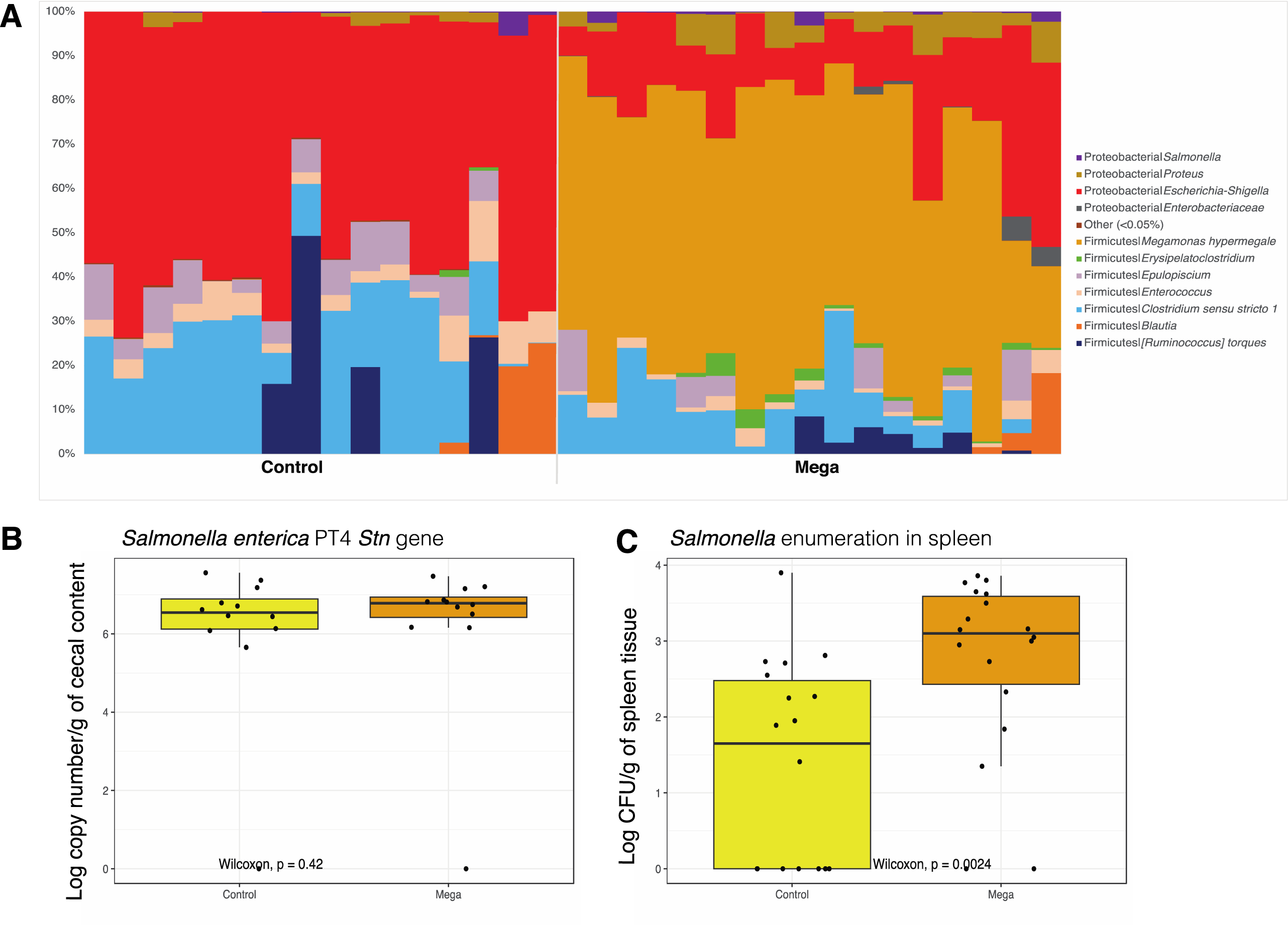
**(A)** Bar plots showing the relative abundances of bacterial species detected in cecal samples obtained from 14-day-old chicks from Control and Mega treatments in EXP2. Dendrograms showing the effect of treatments on *Salmonella* colonization in **(B)** cecal contents based on *Stn* gene quantification by qPCR and in **(C)** spleen tissues based on culturing and enumeration method.

In EXP3, differential abundance analysis indicated that DC and DC + Mega birds presented higher relative abundance of *B. gallinaceum*, *B. uniformis*, *A. finegoldii*, *P. vulgatus*, and *B. viscericola*, and lower relative abundance of *Escherichia-Shigella*, *Proteus*, *Enterococcus*, *Clostridium sensu stricto 1*, and *L. crispatus* than birds in the Control group. In addition, DC + Mega birds showed a lower relative abundance of *Salmonella*, and higher levels of *F. massiliensis*, *R. torques*, and *M. hypermegale* than Control birds. Comparisons between DC and DC + Mega treatments indicated DC cecal microbiota to be enriched for *Escherichia-Shigella* and *L. agilis*, whereas the cecal microbiota of DC + Mega showed enriched *M. hypermegale*, *Clostridium sensu stricto 1*, *R. torques*, and *F. massiliensis* (Figure 5). While 16S sequencing results indicated a reduction in *Salmonella* load in DC + Mega treated birds compared to control group (*p*<0.05), no differences in *Salmonella* load were observed using qPCR (*p* = 0.76). However, the mean log_10_ copy number/g of cecal content of *stn* gene was numerically lower in DC and DC + Mega treated birds compared to control (Figure 6A). *Salmonella* load in the spleen was not different (*p* = 0.08) with *Salmonella* being undetectable in most samples (Figure 6B). Correlation analysis performed between taxa present in the inocula and the baseline microbiota indicated *M. hypermegale* to be negatively correlated with *Escherichia*/*Shigella*, whereas *Salmonella* was negatively correlated with *B. viscericola* and positively associated with *P. vulgatus* and *A. finegoldii*.

**Figure 5.**
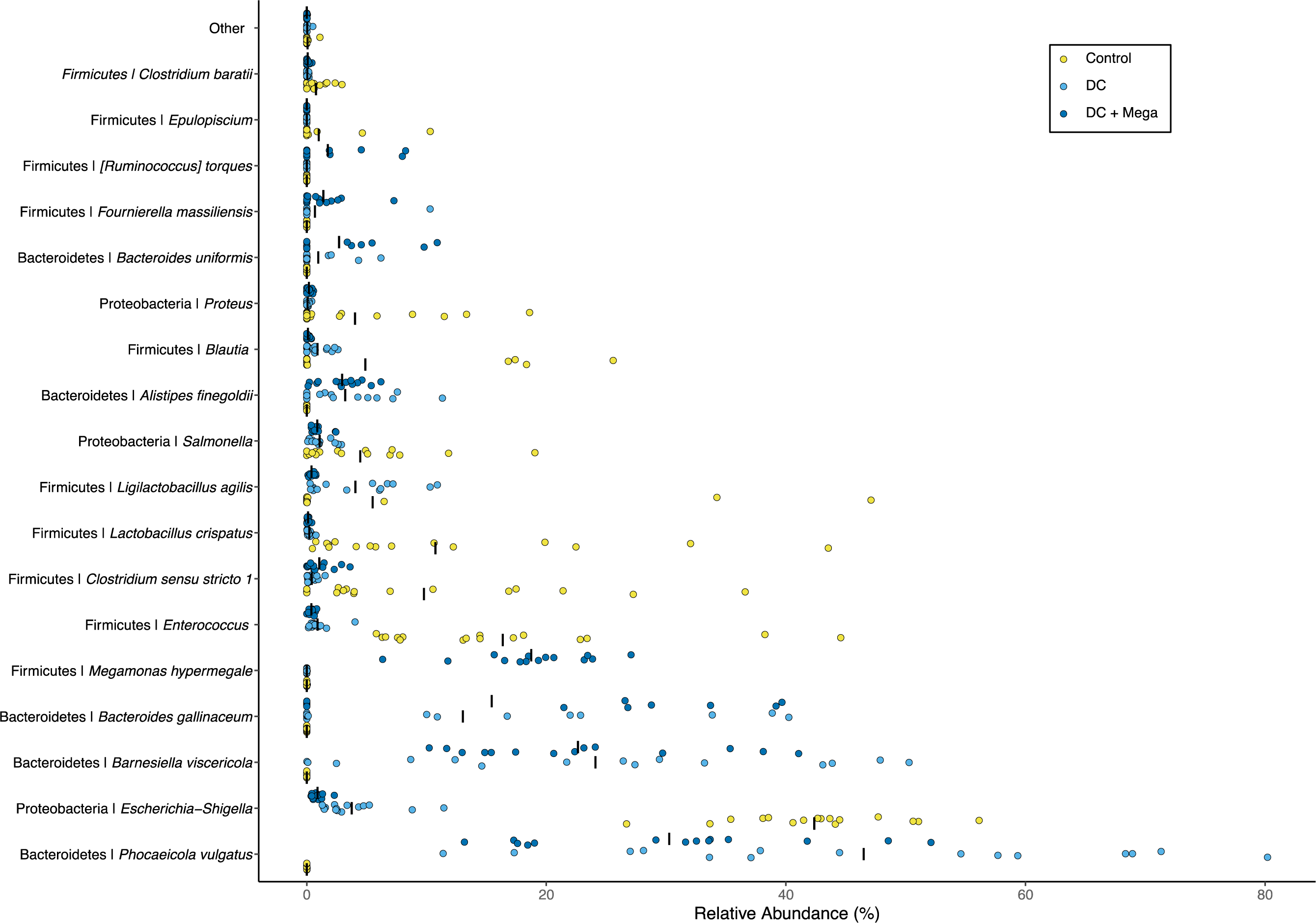
The relative abundance of microbial taxa that were shown to be differentially abundant in the cecal microbiota of 14-day- old broilers from Control, DC, and DC + Mega treatments in EXP3 according to DESeq2 analysis. Dots represent the relative abundance of taxa in individual samples, and bars represent the average relative abundance in each treatment. The relative abundance of taxa that were not differently abundant is shown as “Other”.

**Figure 6.**
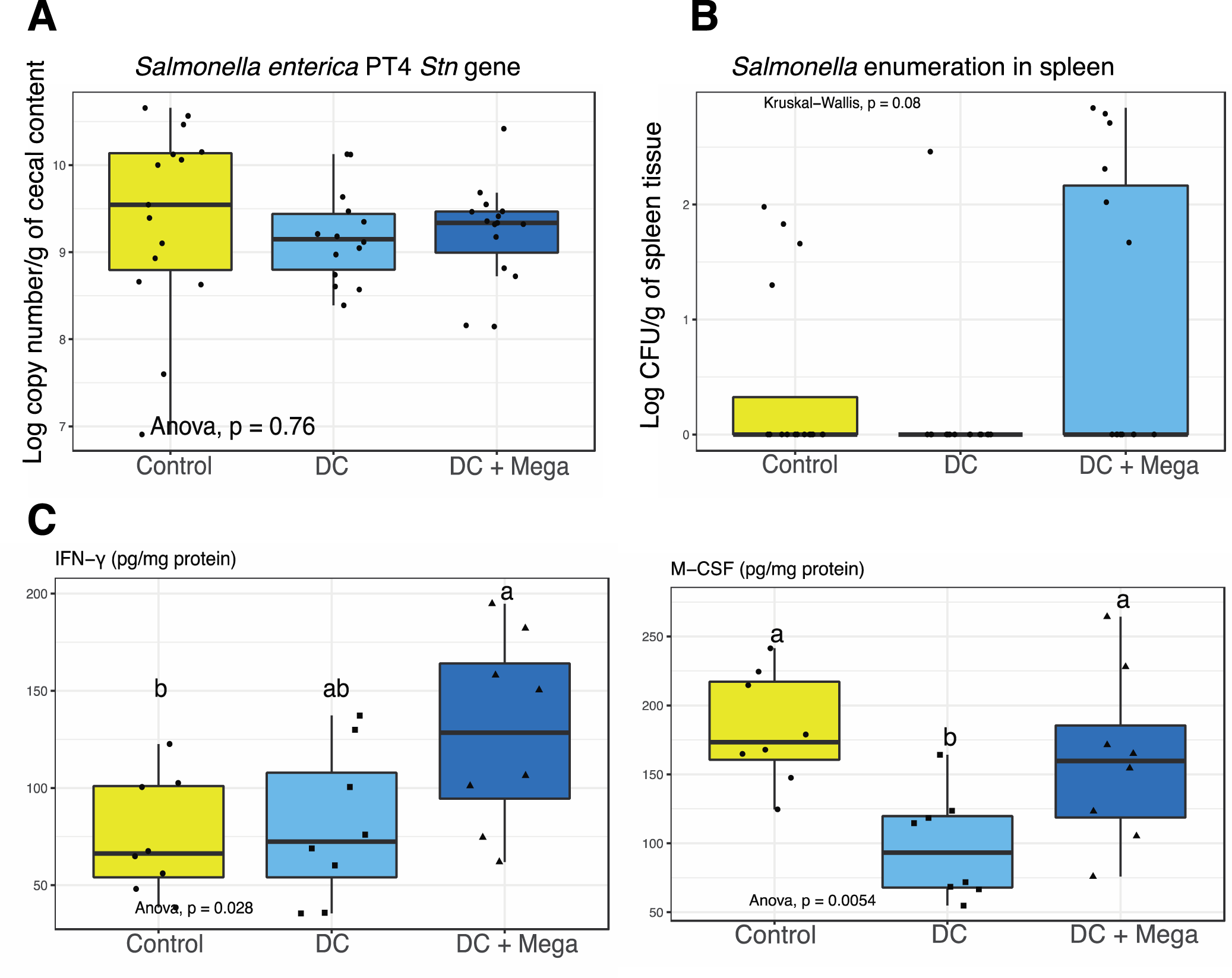
Boxplots showing the effect of Control, DC and DC + Mega treatments in EXP3 on *Salmonella* colonization in **(A)** cecal contents based on *Stn* gene quantification by qPCR and in **(B)** spleen tissues based on culturing and enumeration methods. (**C)** Boxplots showing the effects of treatments on the concentration of IFN-γ and M-CSF in the cecal tissues of 14-day-old broilers. Superscripts with different letters indicate significant differences at α = 0.05.

Birds in the DC + Mega treatment showed a higher level of IFN-γ (*p* = 0.028) in cecal tissue than birds in the Control group. Cecal tissue of birds in the DC treatment exhibited the lowest M-CSF concentration among treatment groups (*p* = 0.005) (Figure 6C), which may coincide with the lower relative abundance of *Salmonella* and the pattern of lower spleen translocation in this group. Cecal concentration of isovalerate was higher in birds from DC and DC + Mega treatments compared with Control. Valerate concentration was higher in DC + Mega compared to Control birds (*p* = 0.045). Propionate (*p* < 0.001) concentration was higher in DC + Mega treated birds compared with birds in Control and DC treatments (Figure 7).

**Figure 7.**
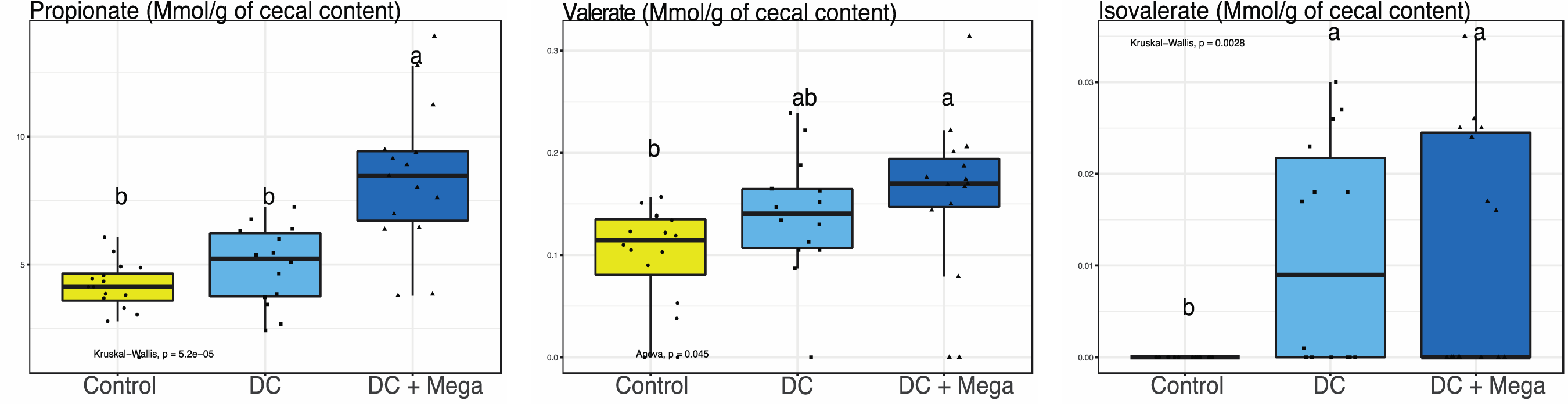
Boxplots showing the effect of Control, DC, and DC + Mega treatments in EXP3 on the concentration of short-chain fatty acids in the cecal content of 14-day-old broilers. Superscripts with different letters indicate significant differences at α = 0.05.

### B. viscericola and P. vulgatus effectively colonize the chicken ceca

Colonization ability and efficiency of bacteria in the inocula were determined based on the prevalence of isolates in the ceca of inoculated birds as well as on the ratio between the number of reads in the cecal samples and in the inocula. *A. finegoldii*, *B. gallinaceum*, *B. viscericola*, *P. vulgatus*, *L. crispatus*, and *L. agilis* were detected in more than half of the inoculated birds and demonstrated high colonization ability. Particularly, *B. viscericola*, *P. vulgatus* and *L. agilis* were detected in all birds. The ratio between the average number of reads in samples and in inocula was higher than 1 for *B. viscericola* and *P. vulgatus* which were considered as highly efficient colonizers. Although *B. uniformis* was not detected in most of the inoculated birds, once introduced, the observed reads in inoculated birds were high, thus this isolate was considered as having poor colonization ability, but to be an efficient colonizer once introduced. *M. hypermegale* failed to colonize the ceca when introduced from a frozen glycerol stock in EXP1 but colonized all the birds and presented a high abundance when introduced as fresh culture in EXP2 and EXP3. *B. mediterraneensis*, *L. aviarius*, and *S. variabile* consistently failed to colonize the chicken gut (Figure 8).

**Figure 8.**
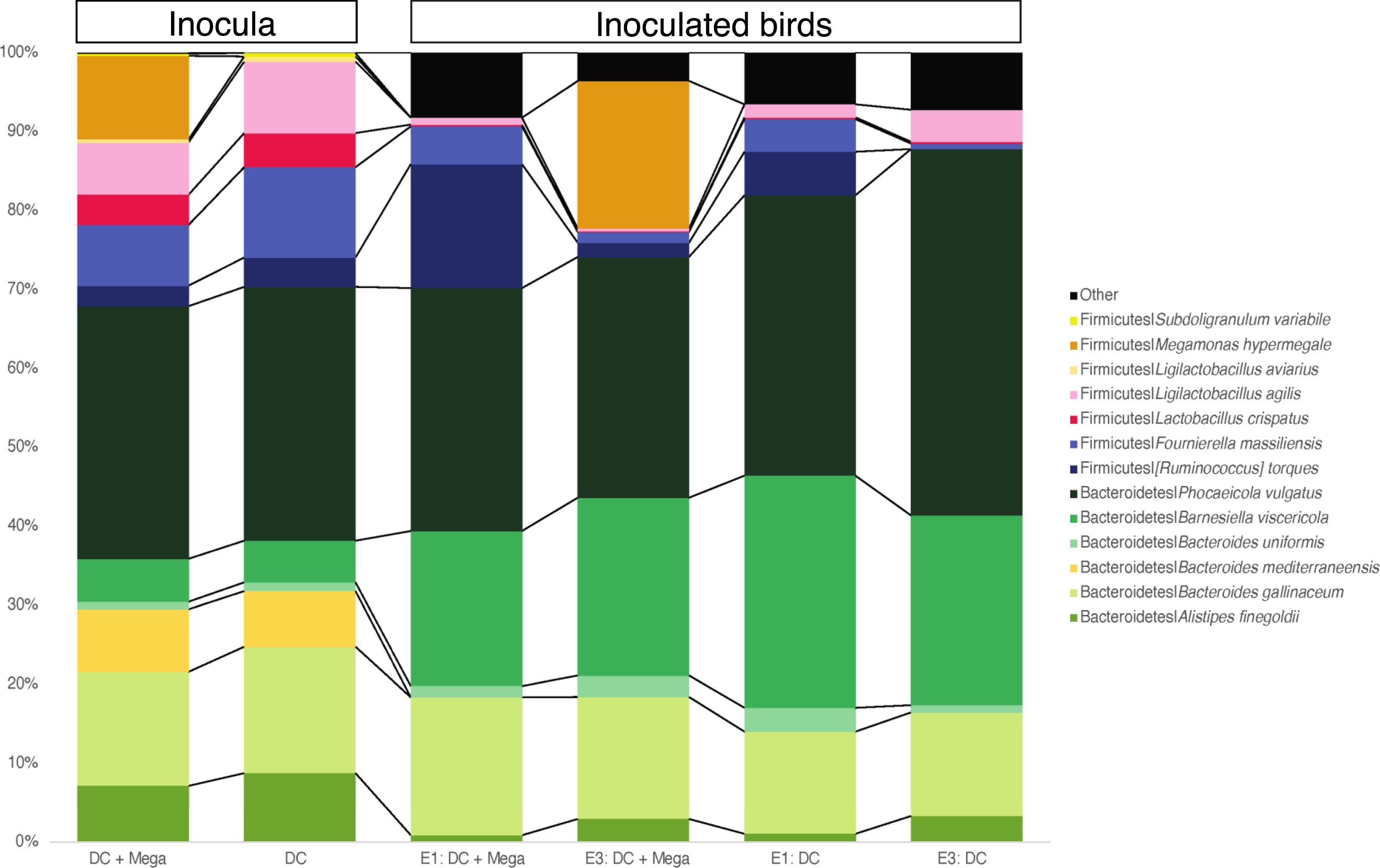
Barplots showing the relative abundance of species included in the DC + Mega and DC inocula and the average relative abundance of these species in the inoculated birds in EXP1 and EXP3. Species present in the birds but not introduced by the inocula were combined as “Other” and are shown in black.

## DISCUSSION

In this study, a DC containing 12 bacterial isolates harvested from adult chicken ceca was inoculated to young chicks, resulting in increased phylogenetic diversity and substantial changes in the cecal microbiota composition evaluated at 14 days old in the context of a *Salmonella* challenge. Strikingly, Bacteroidetes phylum made up more than 75% of the cecal microbiota of DC-inoculated birds but was completely absent in Control birds. Bacteroidetes species were reported to be host-adapted (Kollarcikova et al., 2020) and *Alistipes*, *Bacteroides*, *Barnesiella*, and *Phocaeicola* genera were found to be good colonizers of the chicken ceca, either when introduced as isolates or as part of complex and defined communities (Glendinning et al., 2022; Kollarcikova et al., 2020; Kubasova et al., 2019; Poudel et al., 2022). Bacteroidetes are non-spore forming and sensitive to oxygen, therefore, they have low ability to survive in the environment and are likely to be lost, reduced, or to colonize with delay in broilers due to practices employed in the poultry industry that hinder the contact between newly hatched chicks and their parental microbiota (Karasova et al., 2022; Nurmi & Rantala, 1973). Consequently, it is likely that, once Bacteroidetes are introduced to the gut environment, they can occupy available niches and efficiently engraft.

Despite the substantial differences in cecal microbiota composition, the differences in host responses measured in DC-inoculated and Control birds were subtle. Changes were observed in IFN-γ, M-CSF, propionate, isovalerate and valerate concentration in the ceca, but no significant differences were found in body weight, small intestine morphology and other cytokines measured. Similarly, a previous study found that the inoculation of nine bacteria obtained from chickens affected the chick gut microbiota but caused only a transient increase in systemic IgA levels (Zenner et al., 2021).

Birds colonized with DC + Mega had higher concentrations of valerate and isovalerate compared to control birds, and higher concentration of propionate compared to both Control and DC inoculated birds. Consistent with the current study, we previously found that the concentrations of valerate and propionate were higher in cecal contents of chicks inoculated with microbial cultures and cecal contents from adult birds (Marcolla et al., 2023b). This suggests that inoculation with the DC community partially recapitulated the effects of introducing a complex microbial community regarding the production of SCFAs, with *M. hypermegale* likely playing a major role in propionate production. *M. hypermegale* is a propionate producer and metabolizes free H2 favouring the production of SCFAs by other members of the community (Rychlik, 2020; Sergeant et al., 2014). Although the introduction of DC or DC + Mega were shown to reduce the relative abundance of *Salmonella* in the cecal contents as indicated by 16S rRNA sequencing, only a numeric decrease in *stn* gene abundance was found using qPCR analysis.

We speculated two main reasons why the inoculation with DC did not significantly impact *Salmonella* cecal load compared with Control birds. First, the DC we inoculated had a limited number of microorganisms. Significant reductions in *Salmonella* load were usually observed in studies inoculating defined communities containing 25 species or more (Gleeson et al., 1989; Impey et al., 1982c; Stavric et al., 1985), or using commercial products (Kerr et al., 2013) from which the bacterial composition is not disclosed but is likely to include diverse species. As an exception, a study found that inoculation of germ-free chicks with ten bacterial isolates harvested from the ceca of feral chickens resulted in a significant reduction in *Salmonella* load and ameliorated intestinal inflammation (Wongkuna et al., 2024). In that way, rather than a low number of microorganisms, it is possible that the microorganisms included in our study may not be highly effective to promote *Salmonella* resistance. A second possible explanation is the fact that the baseline microbiota of our birds was enriched with *Escherichia/Shigella,* which made up an average of 42.0 ± 12.0 % of the cecal microbial community in Control birds; however, in birds inoculated with DC and DC + Mega, the average relative abundance of *Escherichia/Shigella* was less than 5%. It is possible that the high abundance of *Escherichia/Shigella* within the ceca of Control birds promoted a similar level of protection of hosts to *Salmonella* infection to that of the DC or DC + Mega treatment. Commensal *Enterobacteriaceae* species were previously demonstrated to protect chickens against *Salmonella* infection through oxygen competition (Litvak et al., 2019). The proposed mechanism indicates that, in the presence of inflammatory signalling, butyrate produced by clostridia stimulate peroxisome proliferator-activated receptor gamma signalling pathway that results in mitochondrial oxidation and epithelial hypoxia (Litvak et al., 2019). The maintenance of epithelial hypoxia by clostridia and the consumption of oxygen by *Enterobacteriaceae* hinder the ability of *Salmonella* to perform aerobic respiration with decreased colonization ability (Litvak et al., 2019). In the present study, *Clostridium Stricto Sensu 1* made up an average of 16.1% of the microbial community in the Control birds, but had lower relative abundance in DC and DC + Mega groups (0.8% and 2.3%, respectively). Despite the absence of differences in *Salmonella* load quantified by qPCR, the fact that DC and DC + Mega reduced the relative abundance of *Escherichia/Shigella* and *Salmonella*, as well as reduced *Enterobacteriaceae* load is a positive outcome. Currently, avian pathogenic *Escherichia coli* is one leading cause of antibiotic treatment in broiler flocks in Canada (CIPARS, 2019), and the use of DC could be investigated as a strategy to mitigate the occurrence of *Escherichia/Shigella-*associated diseases.

In EXP1, *M. hypermegale* provided from a frozen glycerol stock failed to colonize the birds. This was largely unexpected, as *Megamonas* has been shown to be a good colonizer in birds inoculated with frozen cecal contents, cecal cultures, and competitive exclusion products (Dame-Korevaar et al., 2020; Marcolla et al., 2023b). Even if accidentally exposed to oxygen during the inoculation procedure, the *M. hypermegale* strain used in our study was shown to survive for at least 30 minutes. Therefore, it is more likely that the *M. hypermegale* glycerol stock was affected by the freeze-thaw process, since fresh *M. hypermegale* provided at a low dose successfully colonized the ceca of inoculated birds. Freeze-thaw process could also have affected the other three species that failed to colonize (*B. mediterraneensis*, *S. variabile*, and *L. aviarius*). In EXP2, birds were inoculated with fresh *M. hypermegale* broth, resulting in successful colonization, however, birds in Mega treatment had higher *Salmonella* counts in the spleen than Control birds, indicating that introduction of *M. hypermegale* alone, without and accompanying DC, might be detrimental.

Despite the observed effective colonization rate of *M. hypermegale in vivo, in vitro* acid tolerance assay indicated that pH 2 or 3 killed *M. hypermegale*, which is somewhat contradictory. This difference could be due to the 3-h incubation period used in the *in vitro* assay being longer than the retention time within the gizzard and proventriculus *in vivo*. Nine-day-old chickens have retention times ranging from 55 to 480 minutes (Rougière & Carré, 2010), and therefore, future *in vitro* acid tolerance assays should be optimized to mimic *in vivo* conditions accordingly. In addition, the presence of digesta can promote increased acid tolerance by buffering pH and providing nutrients, such as glucose, which can improve bacterial survival (Corcoran et al., 2005). Nonetheless, given *M. hypermegale* provided as a fresh broth culture colonized all inoculated birds at relatively high rates, it is safe to state that it presents sufficient resistance to acidic conditions. The two *in vitro* inhibition assays targeting *Salmonella* resulted in different outcomes: while co-culture assay indicated inhibitory effect of *Salmonella* by *M. hypermegale*, the same was not observed using the slab method. These phenotypes might be explained by differences in metabolism of bacteria when growing in solid or liquid media (Michael & Waturangi, 2023) and optimization of *in vitro* methods is necessary to better understand such differences.

Overall, the results indicated that inoculation with DC and *M. hypermegale* can cause significant shifts in microbiota composition without major effects on broiler physiology or the ability to inhibit *Salmonella* colonization. Among the limitations of this study, the load of each species included in the DC was not quantified. Thus, it is possible that *B. mediterraneensis*, *L. aviarius*, and *S. variabile*, which consistently failed to colonize the birds, were not provided at sufficient load to promote colonization. On the other hand, it is possible that species deemed efficient colonizers were favored by being provided at higher loads. Future studies are warranted using individual isolates introduced at known amounts for a fair comparison of colonization ability. In addition, a comparison between the introduction of isolates using fresh and frozen cultures is needed, given the observed inability of *M. hypermegale* to engraft when provided as a frozen stock, which contrasted with a high colonization ability when provided as a fresh culture.

The DC was designed aiming to promote the establishment of a microbiota community resembling that of a typical broiler, and included the same number of species from Bacteroidetes and Firmicutes phyla (6 each), because major differences in the relative abundances of these phyla were observed between intensive and extensive rearing systems (Marcolla et al., 2023b). Proteobacteria species were not included in the DC as they are highly abundant in the microbiota of newly hatched chicks. According to previous data, a typical broiler microbiota would present a relative abundance of Bacteroidetes and Firmicutes ranging from 20% to 65%, and that of Proteobacteria ranging from 2% to 10%. Despite the presence of Firmicutes in the DC and of Proteobacteria in the baseline community, the microbiota of DC-treated birds was dominated by Bacteroidetes, reaching up to 96% in some individuals, which could be reflective of an unbalanced microbial composition. In addition, it is unclear why some of the species included in the DC failed to colonize the birds (*B. mediterraneensis*, *L. aviarius*, and *S. variabile*). Therefore, future studies are warranted using different combinations of isolates at known concentrations to better understand community composition changes in chicks exposed to DC and to promote colonization with a typical broiler microbiota that would aid the evaluation of individual microbial species.

## CONCLUSION

We concluded that the introduction of a DC containing 12 bacterial species to day-old chicks caused major changes in cecal microbiota composition with subtle effects on host physiology and the ability to resist *Salmonella* challenge. We identified *A. finegoldii*, *B. gallinaceum*, *B. viscericola*, *P. vulgatus* and *L. agilis* as effective colonizers of the chicken gut and found that the introduction of the DC causes significant reduction in the relative abundance of *Escherichia-Shigella* and *Salmonella*, as well as *Enterobacteriaceae* load, which could potentially be explored as a strategy to control the occurrence of *Escherichia/Shigella*-associated diseases.

## ACKNOWLEDGMENTS

This project was funded by Alberta Agriculture and Forestry and a Natural Sciences and Engineering Discovery Grant (RGPIN-2019-06336) held by B.P.W. C.S.M. was supported by Alberta Innovates and the Dr. Michael E. Stiles Graduate Scholarship in Applied Microbiology.

B.P.W. was supported by the Canada Research Chairs program. The funders had no role in study design, data analysis, or interpretation.

B.P.W. obtained funding for the research and contributed to the study design and oversight;

C.S.M. designed the experiments, performed sample collection and analysis, analyzed data, interpreted results, prepared figures, and drafted manuscript. T.J. performed sample collection and analysis, analyzed data, and interpreted results. K.T. conducted *in vitro* experiments. U.S.S and L. U. contributed to laboratory analysis.

B.P.W., C.S.M., and T.J. edited and revised the manuscript. All authors approved final version of manuscript.

